# Neuro-behavioral impact of Tourette-related striatal disinhibition in rats

**DOI:** 10.1101/2025.02.02.636049

**Authors:** Joanna Loayza, Charlotte Taylor, Jacco Renstrom, Rachel Grasmeder Allen, Stephen Jackson, Tobias Bast

## Abstract

Tourette syndrome has been linked to reduced GABAergic inhibition, so called neural disinhibition, in the dorsal striatum. Dorsal-striatal neural disinhibition in animal models, caused by local microinfusion of GABA-A-receptor antagonists, produces striatal local field potential (LFP) spike-wave discharges and tic-like movements resembling motor tics. Here, we characterized further the neuro-behavioral impact of striatal disinhibition, by unilateral picrotoxin infusion into the anterior dorsal striatum, in adult male Lister hooded rats. In vivo electrophysiology under anesthesia revealed enhanced neuronal burst firing in the striatum, alongside spike-wave LFP discharges. In freely moving rats, striatal picrotoxin reliably induced tic-like movements, which mainly involved lifting of the contralateral forelimb and concomitant rotational movements of head and torso, as well as occasional rotations of the whole body around its long axis. Prepulse inhibition (PPI) of the acoustic startle response was not affected, but startle reactivity tended to be reduced. Both locomotor and non-ambulatory movements, measured in photo-beam cages, were increased. Our findings suggest that, apart from generating tic-like movements, dorsal striatal disinhibition may contribute to hyperactivity, which is often comorbid with Tourette syndrome. Enhanced striatal burst firing may be important for these behavioral effects. Our findings do not support that striatal disinhibition contributes to PPI disruption, which has been associated with Tourette syndrome and suggested to contribute to tic generation.

**Significance Statement:** Striatal disinhibition has been implicated in tic generation. Using a rat model, we show that a key neural effect of striatal disinhibition is enhanced firing of striatal neurons in bursts. We also characterized further the tic-like movements caused by unilateral disinhibition of the anterior dorsal striatum. These mainly involved lifting of the contralateral forelimb and concomitant rotational movements of head and torso, as well as occasional rotations of the whole body around its long axis. Striatal disinhibition did not affect prepulse inhibition (PPI) but caused locomotor hyperactivity. This suggests that striatal disinhibition may contribute to the general hyperactivity often comorbid with Tourette syndrome, but that other brain mechanisms underlie the PPI deficits that have been reported in the condition.

## 1. Introduction

Loss of GABAergic inhibitory interneurons has been reported in the striatum of patients with Tourette syndrome (TS) (Brady et al., 2022; Kalanithi et al., 2005; Kataoka et al., 2010; Lennington et al., 2016) and striatal disinhibition is suggested to play a key role in causing tics (Albin & Mink, 2006; Jackson et al., 2015; Kurvits et al., 2020; Tremblay et al., 2015). In support of this, dorsal-striatal microinfusion of GABA-A receptor antagonists like picrotoxin and bicuculline cause tic-like movements in rodents (Bronfeld et al., 2013; Klaus & Plenz, 2016; Pogorelov et al., 2015) and non-human primates (McCairn et al., 2009; Worbe et al., 2009).

Electrophysiological recordings in rats and non-human primates show that striatal disinhibition causes large amplitude local field potential (LFP) spike-wave discharges, alongside increased neuronal firing within the striatum (Darbin & Wichmann, 2008; Israelashvili & Bar-Gad, 2015; Jayasinghe et al., 2017; Klaus & Plenz, 2016; McCairn & Isoda, 2013; McKenzie & Viik, 1975; Muramatsu et al., 1990; Tarsy et al., 1978; Vinner Harduf et al., 2021). Disinhibition-induced striatal LFP spike-waves were associated with tic-like movements in awake animals (Israelashvili & Bar-Gad, 2015; McCairn et al., 2009) but also persisted during sleep when tic-like movements were markedly reduced, suggesting a partial dissociation from the expression of tic-like movements (Vinner Harduf et al., 2021). In the medial prefrontal cortex (mPFC) and hippocampus, neural disinhibition particularly enhanced neural firing in ‘bursts’ (Bast et al., 2017; McGarrity et al., 2017; Pezze et al., 2014). Bursts are periods of relatively high neuronal firing that are separated by periods of comparatively little firing and are particularly effective in driving the targets of neuronal projections (Cooper, 2002; Izhikevich et al., 2003; Lisman, 1997). Therefore, regional neural disinhibition may drive brain-wide activation through enhanced burst firing. However, the impact of striatal disinhibition on burst firing within the striatum remains to be examined.

TS is often co-morbid with hyperactivity (Freeman et al., 2000; Sheppard et al., 1999) and has been associated with impaired sensorimotor gating, as reflected by reduced prepulse inhibition (PPI) (Castellanos et al., 1996; Swerdlow et al., 1994, 2001). Based on the latter, Swerdlow and colleagues proposed that tics are due to failed automatic gating of sensory information (which is experienced in the form of premonitory urges in TS patients) (Swerdlow, 2013; Swerdlow & Sutherland, 2005). Supporting a link between tics and impaired sensorimotor gating comes from transgenic mouse models of TS that showed both tic-like movements and PPI deficits (Castellan Baldan et al., 2014; Godar et al., 2016). The striatum, including the dorsal striatum, has been suggested to be a key region in the modulation of PPI (Baldan Ramsey et al., 2011; Kodsi & Swerdlow, 1995), but the impact of striatal disinhibition on PPI remains to be examined. Moreover, many studies in animals with striatal disinhibition reported informal observations of locomotor hyperactivity, alongside tic-like movements (which may have been referred to as ‘myoclonic jerks’, ‘forelimb tremor’, or ‘stereotyped sniffing’) (Bronfeld et al., 2013; Costall & Naylor, 1975; Israelashvili et al., 2020; Morgenstern et al., 1984; Tarsy et al., 1978; Worbe et al., 2009; Yael et al., 2016; Yoshida et al., 1991), although locomotor measurements following striatal disinhibition, compared to an appropriate control manipulation, have not yet been reported.

Here, we first aimed to confirm and extend previous in vivo electrophysiological studies in the rat striatal-disinhibition model by examining the impact of picrotoxin infusion into the anterior dorso-lateral striatum on striatal LFP spike-wave discharges and multi-unit activity, including burst parameters, in anesthetized rats (Study 1). Second, we characterized tic-like movements in freely moving rats caused by unilateral anterior dorsal striatal disinhibition by picrotoxin, including their time course and key features (Study 2). Third, we examined the impact of such striatal disinhibition on prepulse inhibition (PPI) of the acoustic startle response and on locomotor activity (Study 3).

## 2. Materials and Methods

### 2.1. Rats

In total 50 male adult Lister hooded rats weighing between 290 g and 360 g and approximately 10-11 weeks old at the time of surgery were used (Study 1: 26 rats, Charles River or Envigo, UK; Study 2: 8 rats, Charles Rivers, UK; Study 3: 16 rats, Envigo, UK). For final sample sizes, exclusion criteria and sample size juslficalon, see seclon 2.7 ‘Experimental design’. Rats were housed in groups of four in two-level high-top cages (462 mm x 403 mm x 404 mm; Techniplast, UK), with an alternating light/dark cycle of 12 h per phase (lights were on between 7 am and 7 pm), at a temperature of 21±1.5 °C and a humidity of 50±8%. Food (Teklad global 18% protein diet 2019, Harlan, UK) and water were provided ad libitum. All experimental procedures were carried out during the light phase. Prior to the start of any procedures, rats were handled daily to habituate them to the experimenters. All procedures were conducted in accordance with the requirements of the United Kingdom (UK) Animals (Scientific Procedures) Act 1986 under Home Office Project Licenses 30/3357 and PP1257468. Experimental methods are reported in line with the ARRIVE guidelines (Lilley et al., 2020).

### 2.2. Implantation of infusion cannula-microwire recording array assembly (Study 1) and guide cannula (Studies 2 and 3) into the anterior dorsal striatum

Rats underwent stereotaxic implantation of an infusion cannula-microwire recording array assembly (for electrophysiological Study 1) or of an infusion guide cannula (for behavioral studies, Study 2 and 3) into the right anterior dorsal striatum. The target coordinates for the infusion site were +1.5 mm anterior to bregma, +2.5 mm right from the midline, and -5.0 mm ventral from the skull (or -4.5 mm from dura respectively). These coordinates were chosen using ‘Ratlas-LH’, an MR atlas of the male Lister hooded rat brain (Prior et al., 2021) and were adapted from previous studies using striatal infusions of GABA-A-receptor antagonists to induce tic-like forelimb movements in other rat strains (Bronfeld et al., 2013; Klaus & Plenz, 2016). Rats were anesthetized with isoflurane in oxygen (induced with 3-4% and maintained at 1.5–3%; flow rate 1 L/min) and, in Studies 2 and 3, were subcutaneously injected with an analgesic (Rimadyl large animal solution, Zoetis, UK; 1:9 dilution, 0.1 ml/100 g) and with an antibiotic (Synulox, 14% Amoxicillin, Zoetis, UK; 0.2 ml/100 g, s.c.) as prophylaxis against infection that can be a complication with indwelling cannulae. Study 1 was under terminal anesthesia and, therefore, rats did not receive analgesic or antibiotic. Rats were positioned in a stereotaxic frame. EMLA cream, containing a local anesthetic (lidocaine/prilocaine, 5%, AstraZeneca, UK), was applied onto the ear bars to reduce pain, and an eye gel (Lubrithal, Dechra, Denmark) was used to prevent ocular drying. An incision was made in the scalp, the skull was exposed and bregma and lambda were aligned horizontally.

In Study 1, a custom-made assembly of a 33-gauge stainless-steel infusion cannula and a 1×4 or 2×4 microwire-electrode recording array (NB Labs, USA; MicroProbes, USA; or Plexon, USA) was used, similar to our previous studies (McGarrity et al., 2017; Pezze et al., 2014). The arrays consisted of a stainless-steel ground wire and 50 µm Teflon-coated stainless-steel wires, which were about 0.5 mm apart from each other. The infusion cannula tip was positioned approximateley 0.5 mm above the tips of the central electrodes (**Figure 1a**). To implant the infusion cannula-recording array assembly, the bone above the right anterior dorsal striatum was removed with a drill, and the exposed brain was kept moist with 0.9% saline. The end of the cannula was connected to a 1 µl Hamilton syringe via flexible Teflon tubing (0.65 mm OD x 0.12 mm ID, Bioanalytical Systems, Inc). Before the Hamilton syringe was connected to the Teflon tubing, the tubing was filled with 0.75 µl distilled water and 0.25 µl air. Once connected to the tubing, the syringe was then pushed to 0.5 µl, before being pulled back to 0.75 µl. This method created one 0.25 µl air bubble at the end of the infusion cannula tip, to reduce the risk of leakage and drug diffusion before the infusion, and another bubble where the syringe was connected to the tubing. The latter bubble was monitored for movement during the infusion to verify that the intended infusion volume moved through the cannula. Infusion cannula and tubing were filled with the infusion solution (picrotoxin or saline). The assembly was fixed to the arm of the stereotaxic frame, such that the electrode array was perpendicular to the midline of the brain and anterior to the infusion cannula. The infusion-recording assembly was slowly lowered until the infusion cannula tip reached the target coordinates in the striatum. Upon reaching the target coordinates, there was at least a 30 min stabilisation period before recording started. Throughout the electrophysiological recording, rectal temperature was measured using a rectal probe and maintained at about 37 °C, using a homeothermic blanket control unit (Harvard Apparatus Ltd, USA).

**Figure 1:**
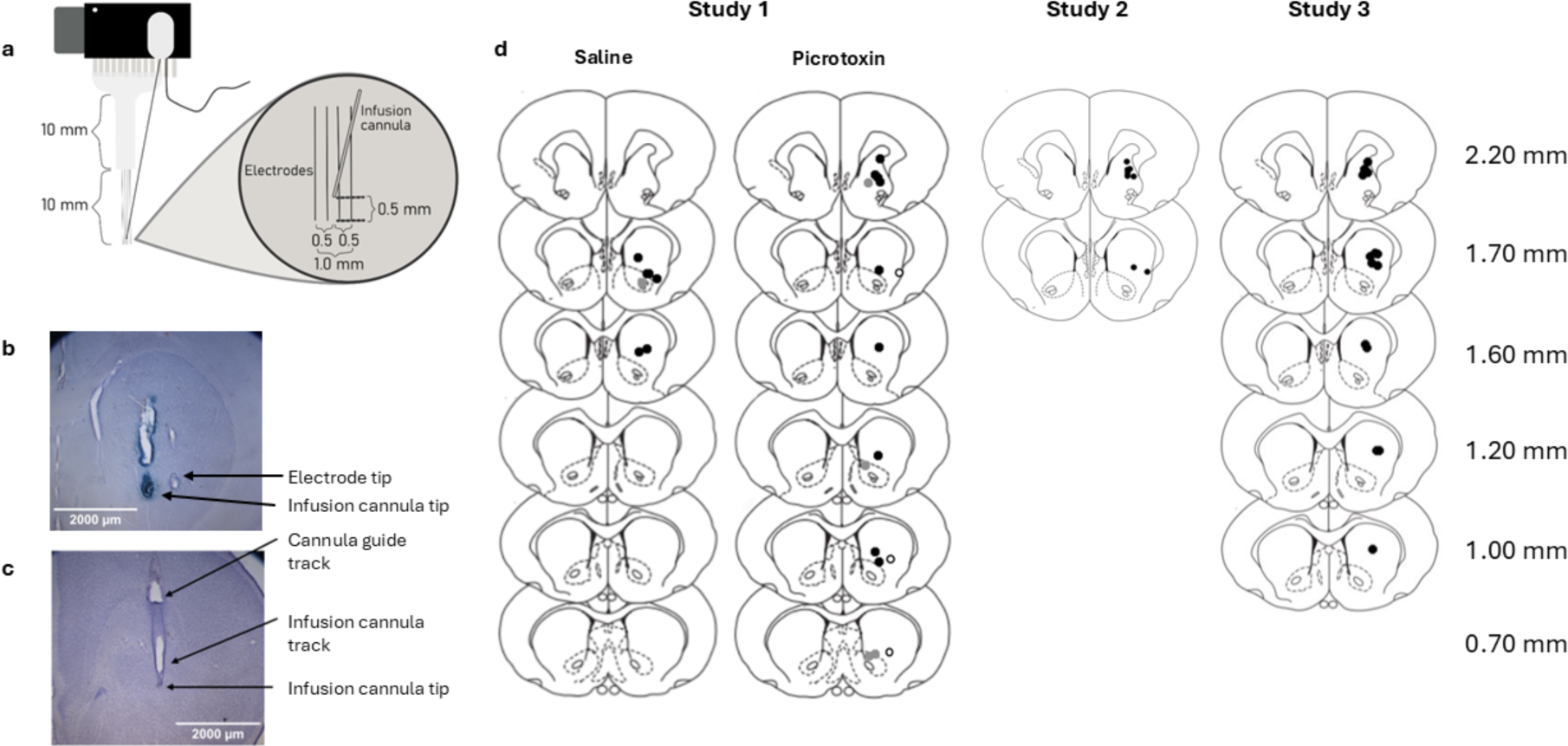
Electrode and infusion cannula placements. *Note.* (a) Sketch of the assembly of infusion cannula and 2×4 microwire-electrode array. The array was arranged perpendicular to the midline of the brain, with the infusion cannula located in the center of the array, about 0.5 mm above the electrode tips. (b) Coronal section of the striatum from a rat used for in vivo electrophysiological measurements; the section shows electrolytic lesions marking the location of the infusion cannula (left) and of the most lateral electrode (right). (c) Coronal section of the striatum from a rat used in the behavioral studies; the section shows tracks left by the guide and infusion cannula. (d) Approximate locations of infusion cannula tips in the anterior dorsal striatum for all rats included in the data analysis in Studies 1 to 3. Locations are shown on coronal plates adapted from the atlas by Paxinos and Watson (1998), with numbers indicating distance from bregma in millimeters, as shown in the atlas. For Study 1, placements are shown separately for the two groups that received infusions of saline or picrotoxin. The figures show the placement of the tips of the most medial (grey dot) and lateral (white dot with black outline) electrodes, as well as of the infusion-cannula tips (black dot).

In Studies 2 and 3, a hole was drilled unilaterally above the infusion site and an infusion guide cannula (C315G-SPC guide, 26-gauge, 4.4 mm below pedestal, Plastics One Inc., Bilaney Consultants, UK), with a stainless-steel stylet (C315DC/SPC to fit 4.4 mm 315G with 2.0 mm projection, Plastics One Inc, Bilaney Consultants, UK) was inserted into the brain, to prevent occlusion of the guide cannula. The guide cannula was secured to the skull using dental acrylic (flowable composite; Henry Schein, Germany) and 5 stainless-steel screws. Screws were placed in front, behind and just to the side of the cannula, to ensure stabilization of the unilateral implant. The incision site was then sutured around the implant. Rats received an intraperitoneal injection of saline (1 ml) for rehydration and were monitored while recovering from the anesthesia. In the days following surgery, rats were checked, weighed, and injected with antibiotic (Synulox, 14% Amoxicillin, Zoetis, UK; 0.2 ml/100 g, s.c.), as well as daily handled to habituate rats to the infusion procedure. Rats were given at least 7 days to recover from surgery before commencing the experiment.

### 2.3. Striatal picrotoxin infusion

Picrotoxin (Sigma-Aldrich, UK) was dissolved in saline at 300 ng/0.5 µl, aliquoted and kept frozen (up to 6 months) until use for infusion. Previous studies suggested that unilateral infusion of this dose into the striatum would have robust neural and behavioral effects, without causing seizures or other adverse effects. Klaus and Plenz (2016) infused 0.8-1.5 µl of 1 mM (603 ng/µl) picrotoxin solution into the anterior dorsal striatum and reported marked LFP spike-wave discharges within the striatum and tic-like forelimb movements. In addition, Morgenstern et al. (1984) reported occasional convulsions and marked locomotor hyperactivity with bilateral infusions of picrotoxin at 500 ng/1 µl/side into the ventral striatum, but only locomotor hyperactivity at 250 ng/1 µl/side. In Study 2 we also examined a lower dose, for which we diluted aliquots to 200 ng/0.5 µl in saline.

In Study 1, after baseline neural activity had been recorded for 30 min, the piston of the 1 µl Hamilton syringe was moved slowly (about 0.5 µl/min) to remove the 0.25-µl air plug from the infusion-cannula tip and to inject 0.5 µl of picrotoxin or saline into the striatum. To verify successful infusion, we monitored movement of the air bubble that was trapped where the tubing was connected to the Hamilton syringe. Start and end times of the infusion were noted, so that data recorded during the infusion period could be removed and the start time of the post-infusion period could be identified for the data analysis. After the infusion, recordings continued for a minimum of 60 min.

In Studies 2 and 3, rats were gently restrained to remove stylets and to insert infusion cannulae (C315I/SPC internal 33 gauge to fit 4.4 mm C315G guide cannula with 2 mm projection, Plastics One Inc., Bilaney Consultants, UK) into the guides. The infusion-cannula tip extended 2 mm below the guide into the right anterior dorsal striatum (DV: -5 mm infusion tip from skull). The infusion cannula was connected through polyethylene tubing (PE50 tubing, C313CT), Bilaney Consultants, UK) to a 5 µl syringe mounted on a microinfusion pump. Prior to infusions, the tubing and syringe were filled with distilled water and an air bubble was included before any drug solution was pulled up. Sterile saline (control) (0.9%, 0.5 µl) or picrotoxin (either 300 ng/0.5 µl or 200 ng/0.5 µl in sterile saline) was delivered over 1 min. Movement of the air bubble within the tubing was monitored to verify infusion into the brain. Following infusion, the infusion cannula remained in place for an additional minute to allow absorption of the infusion bolus by the brain tissue. The infusion cannula was then replaced by the stylet and rats were visually inspected for presence of tic-like movements.

### 2.4. Multi-unit and LFP recordings (Study 1)

We recorded extracellular measures of neural activity, using methods similar to our previous studies (McGarrity et al., 2017; Pezze et al., 2014). The electrode array was connected to a multichannel preamplifier (Plexon Inc., USA) via a unity-gain multichannel headstage. The preamplifier amplified the analogue signal from the electrodes by a factor of 1000 and band-pass filtered the signal into multi-unit spikes (250 Hz to 8 kHz) and LFP signals (0.7 Hz to 170 Hz). All recordings were made against ground, with the ground wire of the array clamped to the ear bars of the stereotaxic frame using a crocodile clip and a lead linking the stereotaxic frame to the ground jack on the amplifier. The analogue signals were then processed by the multichannel acquisition processor (MAP) system (Plexon Inc., USA). This allowed for an additional computer-controllable amplification of up to 32,000, further filtering of multi-unit data (500 to 5 kHz), digitization of spikes at 40 kHz (providing 25 µs precision on each channel at 12-bit resolution) and recording of LFP data at 1 kHz.

Multi-unit data and LFP data could be viewed during the experiments, using Real-Time Acquisition System Programs for Unit Timing in Neuroscience (RASPUTIN) software (Plexon Inc., USA). RASPUTIN was used to record neural activity data for the baseline and post-infusion period. LFP data were recorded continuously, and multi-unit spikes were recorded when they exceeded a predefined amplitude threshold of -240 µV.

NeuroExplorer version 4 software (Nex Technolgies, Plexon Inc., USA) was used to calculate firing rate and burst parameters from the multi-unit data and power spectral densities (PSD) from the LFP data, as in our previous study (Pezze et al., 2014). To detect striatal bursts, we used the Poisson surprise method, similar to previous studies (Homayoun et al., 2005; Miller et al., 2008; Pezze et al., 2014; Stevenson et al., 2007). Only bursts with a surprise value of greater than 3 were included in our analysis (Pezze et al., 2014; Stevenson et al., 2007). This value means that there is an approximate probability of 0.05 for similar spike patterns to occur by chance as part of a random spike train. The following burst parameters were calculated for each 5 min block: mean spike frequency, number of bursts, percentage spikes in bursts, mean burst duration, mean spikes in bursts, mean spike frequency in bursts, mean peak spike frequency in bursts, mean interburst interval and mean burst surprise. For the analysis of interburst intervals, one channel (out of 4) had to be excluded in one rat, because this channel did not record more than one burst in one 5 min block, and, therefore, an interburst interval measure was not available for all 5 min blocks. For PSD analysis, the NeuroExplorer software applied fast Fourier transform analysis to the LFP signal. We calculated the Area under the Curve (AUC) of the PSD function from 0.7 - 40 Hz as a measure of overall LFP power for every 5 min block of the pre-infusion and post-infusion recording periods.

### 2.5. Behavioral measurements

#### 2.5.1. Tic-like movements (Study 2)

Rats were placed into one of four rectangular boxes (38 cm x 40 cm x 53 cm, Tontarelli, UK) positioned in a well-lit room (200 lx). An overhead camera (HD PRO Webcam C920, 1080p/30 fps - 720p/30 fps, Logitech, UK) was placed centrally over the boxes, and was used to record the rats’ behavior for subsequent analyses of tic-like movements. A movement was counted as a tic-like movement when it was abrupt and occurred out of the context of the rats’ ongoing behavior. Tic-like movements following picrotoxin infusion into the anterior dorsal striatum in the present study mainly involved lifting of the contralateral forelimb and concomitant rotational movements of head and torso, but some tic-like movements were more intense and involved rotation of the body around the long axis (see Results, Section 3.3.1). The tic-like movements involving whole-body rotations were noted and analyzed both as part of overall tic-like movements and separately. The onset of tic-like movements was determined in reference to the rat being placed in the box. The duration of tic-like movements was calculated by measuring the time between the first and last observed occurrence of a tic-like movement.

#### 2.5.2. Startle and PPI testing (Study 3)

Following striatal infusion, rats were placed immediately into the Startle/PPI test boxes and doors were closed. Startle and PPI was measured using four startle response systems (San Diego Instruments, USA), as in Pezze et al. (2014). Systems were placed in individual sound-attenuating chambers (39 x 38 x 58 cm^3^) that were well lit by individual lights within the chambers (ranging from 228 to 355 W), ventilated and consisted of a clear Perspex cylinder (8.8-cm diameter, 19.5 cm long) on a solid Perspex base connected to an accelerometer, which connected to Reflex Testing software (San Diego Instruments, USA). Once all four rats were placed into their boxes, testing started. Sessions started within 1 to 11 min following infusion (mean = 4.24, SEM = 0.61). The amplitude of the whole-body startle response to an acoustic pulse was defined as the average of 100 x 1 ms accelerometer readings collected from pulse onset.

A test session started with a 5 min acclimatization period of the rat being in the cylinder with a 62 dB(A) background noise level, that continued throughout the session. Following acclimatization, there were three test blocks. (1) The first block consisted of 10 individual startle pulses [40 ms, 120 dB(A) broad-band bursts], allowing for habituation of the startle response to a relatively stable level for the remainder of the session. (2) The second block consisted of a total of 50 trials of 5 different trial types, which were each presented 10 times in a pseudorandom order and with an intertrial interval that varied between 10 to 20 s in duration (average 15 s). The 5 different trial types consisted of pulse-alone trials and four types of prepulse plus pulse trials, in which a weak 20 ms prepulse of 72, 76, 80, 80 or 84 dB(A) preceded the startle pulse by 100 ms. The percentage of PPI induced by each prepulse intensity was calculated as follows: [(mean startle amplitude on pulse-alone trials − mean startle amplitude on prepulse-plus-pulse trials)/(mean startle amplitude on pulse-alone trial)] × 100%. (3) The final block consisted of five startle pulses. Analysis of startle amplitude on pulse-alone trials across all three blocks was conducted to measure startle habituation. The total duration of the test session was 23 min.

#### 2.5.3. Open-field locomotor activity and non-ambulatory movements (Study 3)

Rats were placed in clear Perspex chambers (39.5 cm long x 23.5 cm wide x 24.5 cm deep) with metal grid lids placed in a dimly lit room (light levels in the chambers ranged from 37-60 lx, with the lids off), similar to Pezze et al. (2014). The chambers sat in frames containing two levels of a 4 x 8 photobeam configuration (Photobeam Activity System; San Diego Instruments, USA). As in previous studies (McGarrity et al., 2017; Pezze et al., 2014), locomotor counts were recorded when two successive photobeams within the lower level of photobeams were broken, which indicated that there was movement of the rat’s body from one beam to another, reflecting locomotion. In addition, in the present study, we recorded ‘non-ambulatory movement’ counts when the same beam was interrupted for the second and subsequent times, with no other beam being interrupted in the meantime, which reflected movement of the rat’s body or parts of the rat’s body without locomotion. A 1 s refractory period was used, during which additional interruptions of a beam were not counted, to avoid counting scratching or tail flipping. Additional interruptions of the same beam within the refractory period restarted the refractory period. When a rat blocked multiple beams due to its size, the system counted the last interrupted beam as the ‘currently interrupted’ one. We expected that such non-ambulatory movement counts may correspond partly to tic-like movements, potentially providing an approximate, but, importantly, automated measure of these movements. A session started with the placement of the rat into the center of the chamber and lasted 60 min. For analysis, data was combined into 5 min blocks.

### 2.6. Histology - verification of electrode (Study 1) and infusion cannula placements in the anterior dorsal striatum (Studies 1 to 3)

For Study 1, at the end of each recording, rats were overdosed by increasing the isoflurane levels. Using a lesion marker (Model D.C. LM5A, GRASS Instruments, USA) a constant current (1 mA, 10 s) was passed through the stainless-steel electrodes or infusion cannula. Usually, the most lateral and medial electrode, as well as the infusion cannula, were connected to the plus pole (anode) of the lesion maker (while another electrode was connected to the minus pole) to mark their location and to deposit ferric ions at the tip of the electrode and cannula. The infusion-recording assembly was then removed, and rats were overdosed. Brains were extracted and stored in a 4% paraformaldehyde solution with 4% potassium ferrocyanide for at least 2 days. Following completion of Study 2 and 3, rats were overdosed with sodium pentobarbital (Dolethal, Vetoquinol, UK) and transcardially perfused using 0.9% saline followed by 4% paraformaldehyde solution. Brains were extracted and stored in 4% paraformaldehyde. Brains were sliced into 80 µm thick coronal sections using a vibratome (VT1000 S, Leica, UK) and placed onto microscope slides. Infusion cannula placements were verified using a Leica light microscope (Leica DM1000, camera: Leica DFC295, microscope objective: Leica HCX PL Fluotar 1.25x/0.04). Cannula and electrode placements were mapped onto coronal sections adapted from the rat brain atlas by Paxinos & Watson (1998). Some slides were stained with cresyl violet to take photos, captured with a Zeiss Axioplan light microscope using a QImaging MicroPublishers 5.0 RTV and the Micro Manager software. Images were taken using a 1.25x/0.035NA objective. The images were then white balanced, and a scale bar was applied using the FIJI software.

### 2.7. Experimental design

#### 2.7.1. Study 1

Neural effects of striatal infusion of saline (0.5 µl) or picrotoxin (300 ng/0.5 µl) were compared between-subjects. Based on previous studies demonstrating robust effects of hippocampal and prefrontal picrotoxin infusions on neural activity in the vicinity of the infusion site (McGarrity et al., 2017; Pezze et al., 2014), our target sample size was n=8 rats per group. The experiment was run in multiple batches with the aim to complete successful recordings in the target sample size. In total, there were five batches of rats: eight rats each in the first and second batches, five in the third, one in the fourth, and four in the fifth. For each batch, rats were randomly allocated to a drug group.

The drug allocation was counterbalanced across batches, ensuring similar numbers of rats infused with saline and picrotoxin in different batches. Due to exclusion of rats based on the histological verification of cannula and electrode placements, our saline group included only 6 rats. Rats were excluded due to the following reasons: in 5 rats, the infusion cannula arrays were misplaced; in 2 rats the correct placement of the infusion cannula array in the striatum could not be verified (because we did not find clear electrode or cannula marks); and in 5 rats, infusion could not be verified (because we could not confirm air bubble movement, see Section 2.3). In total, 14 rats (saline, n=6; picrotoxin, n=8) contributed to the electrophysiology data. For 2 rats (infused with saline), data from 2 electrodes were excluded from analysis due to electrode placement outside of the striatum. Analysis was always conducted with a minimum of 2 electrodes per rat.

#### 2.7.2. Study 2

This study used n=8 rats to examine tic-like movements caused by striatal picrotoxin infusion. This sample size was based on previous studies suggesting that striatal disinhibition reliably induced tic-like movements in rodents (Bronfeld et al., 2013; Klaus & Plenz, 2016; Pogorelov et al., 2015). Our expectation was that at least half of the rats would show tic-like movements, and if that had not been the case we would have gradually increased the picrotoxin dose or the infusion volume. Following surgery and recovery, the study lasted 8 days. On day 1, rats were habituated to the observation boxes for 30 min. This was followed by seven additional days, consisting of four infusion days, when rats were monitored for tic-like movements following infusions, separated by washout days to avoid carry-over effects from the preceding infusion day. On infusion days, following a 30 min habituation period in the observation boxes, rats received striatal picrotoxin or saline infusions, before they were placed into the boxes for up to 2.5 h. On the first infusion day (day 2), rats received 300 ng/0.5 µl picrotoxin infusion, to investigate whether this dose caused marked tic-like movements of the contralateral forelimb, without seizures or any other gross adverse side effects. For the second infusion day (day 4), rats were infused with 0.5 µl saline, to verify that saline alone does not cause any behavioral changes and that tic-like movements are caused by striatal disinhibition, rather than nonspecific infusion effects. On the third infusion day (day 6), rats received 300 ng/0.5 µl picrotoxin infusion again, to investigate whether marked tic-like movements can be caused reliably by the same dose within the same rat. On the fourth infusion day (day 8), a lower dose of 200 ng/0.5 µl picrotoxin was used to test if this dose also caused tic-like movements.

#### 2.7.3. Study 3

Study 3 compared the within-subjects effects of striatal picrotoxin (300 ng/0.5 µl) and saline infusions on startle, and PPI, as well as on open-field locomotor activity and non-ambulatory movements. Starting the study with a batch of 16 rats, our target sample size was n=12-16, which would give us a power > 80% to detect differences between striatal disinhibition and striatal saline infusion that correspond to an effect size of Cohen’s d of about 1 or greater, using two-tailed paired t-tests (p < 0.05) (power analysis conducted with GPower 3.1 (Faul et al., 2007)). Following surgery and recovery, rats underwent 5 successive days of startle/PPI testing, and 2 days later, 5 successive days of open-field testing for locomotor activity/non-ambulatory movements. On day 1 of startle/PPI and open-field testing, rats were tested without infusions to habituate them to the testing procedure and to record baseline measurements. On day 2 and 4, rats received one of the two infusions (saline or picrotoxin, with testing order counterbalanced) followed by behavioral testing. To verify that any differences between drug conditions on the infusion day reflected temporary infusion effects, rather than any other confounding factors, rats were re-tested, without any infusion, on the days after infusion (day 3 & 5). Three rats had to be excluded from all or part of the data analysis: two rats had a blocked guide cannula so could not be infused and one rat had to be culled after it showed adverse side effects during the first infusion day of the open-field session following picrotoxin infusion, which included loss of postural control and unresponsiveness, and thus had to be excluded from the open-field data analysis. In total, n=14 rats contributed to the startle/PPI data and n=13 rats contributed to the open-field locomotor/movement data.

### 2.8. Statistical analysis

A significance level of p < 0.05 was accepted for all statistical tests. Analysis was completed using IBM SPSS Statistics 24, and graphs were completed on GraphPad prism (Version 9, GraphPad software, USA).

#### 2.8.1. Study 1, multi-unit and LFP data

Values of the multi-unit and LFP parameters were calculated from the data of each electrode for each 5 min block of the baseline and post-infusion recording periods. For normalization to baseline, values obtained from the individual electrodes were divided by the average values obtained from the same channel during the 6 (5 min) baseline blocks. Normalized values were averaged across all electrodes per individual rat, and these average values were used to calculate the group means used for statistical analysis. Data were examined using ANOVA with drug infusion as the between-subjects variable (saline versus picrotoxin) and time points (5 min blocks) as within-subjects factor. We focused on the interaction infusion x time to assess significant drug infusion effects on the time course of the recording data, as in our previous studies (McGarrity et al., 2017; Pezze et al., 2014). Baseline recordings were compared between infusion groups using independent samples t-tests.

#### 2.8.2. Study 2, tic-like movements

Individual tic-like movements of each rat were counted manually in 5 min blocks, from the video recordings. Using a repeated-measures ANOVA with drug infusion and time point (5 min blocks), data was examined for significant differences between infusions in the number of overall tic-like movements, onset, duration and number of tic-like movements involving whole-body rotations.

#### 2.8.3. Study 3, startle/PPI and locomotor activity/non-ambulatory movement data

Data were analyzed using ANOVA with drug infusion (saline/picrotoxin) and test blocks or trial types (12 x 5 min blocks for open-field testing, three test blocks for startle testing and four different prepulse types for PPI testing) as within-subjects factors.

## 3. Results

### 3.1. Infusion-cannula (Study 1-3) and electrode (Study 1) placements

All rats included in the analysis had their infusion cannula placed within the anterior dorsal striatum (**Figure 1b-d**). Electrode placements in Study 1 (in vivo electrophysiology) were mostly placed within the dorsal striatum, but some medial electrode placements encroached on the dorsal nucleus accumbens (**Figure 1d**, left).

### 3.2. Study 1: Striatal disinhibition caused marked LFP spike-wave discharges and enhanced multi-unit burst firing in the striatum of anesthetized rats

#### 3.2.1. Qualitative description of LFP and multi-unit activity patterns induced by striatal disinhibition

A marked impact of anterior dorsal striatal picrotoxin infusions on multi-unit and LFP recordings in the vicinity of the infusion site in anesthetized rats was evident based on visual inspection of the recordings in most rats (**Figure 2**). In five out of eight rats that received striatal picrotoxin, but in none of the rats that received saline infusions, there were large LFP spike-wave discharges, consisting of a single negative spike followed by a positive wave, and intensified multi-unit bursts during the negative spike. We did not observe LFP characteristics of seizures, including higher-frequency (10-15 Hz) sharp LFP deflections (polyspikes’) superimposed on the large spike-wave discharges (Aupy et al., 2020; Neckelmann et al., 1998; Steriade & Contreras, 1998). Importantly, we also did not observe tic-like movements under anesthesia.

**Figure 2:**
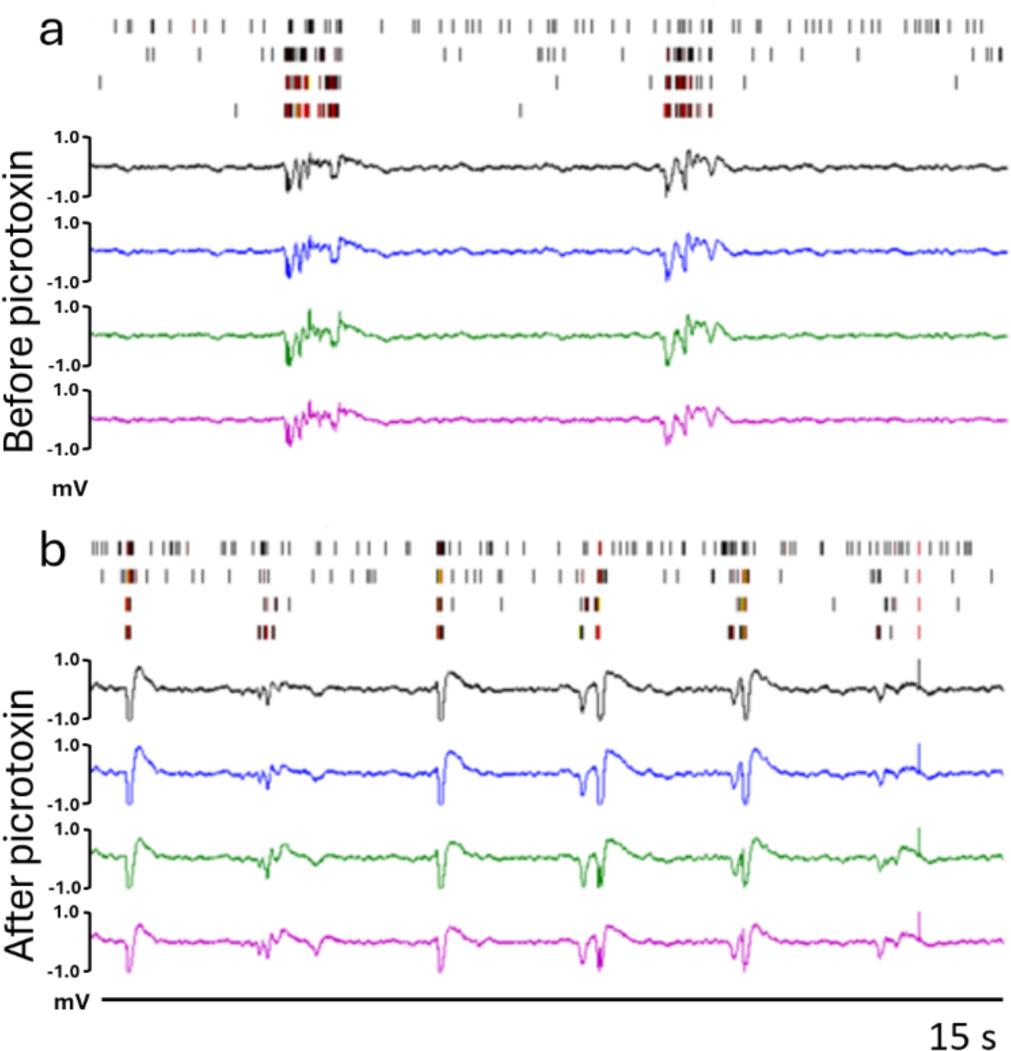
Striatal picrotoxin enhanced LFP spike-wave discharges and multi-unit burst firing in the vicinity of the infusion site. *Note.* Raster plot of multi-unit spike activity (top, electrodes 1-4; black, gold and red vertical lines indicate increasing levels of spiking activity) and LFP traces (bottom, electrodes 1-4) during a 15-s period recorded about 10 min before (A) and after (B) picrotoxin infusion (300 ng/0.5 µl). Following picrotoxin infusion, there were large LFP spike-wave discharges, consisting of a single negative spike followed by a positive wave, and sharp multi-unit bursts during the negative spike. Note that negative LFP spikes were clipped due to their amplitude being out of range.

#### 3.2.2. Mean baseline values of electrophysiological parameters

The mean baseline values of all parameters analyzed (mean spike frequency, number of bursts, percentage spikes in bursts, mean burst duration, mean spikes in bursts, mean spike frequency in bursts, mean peak spike frequency in bursts, mean interburst interval, mean burst surprise and mean overall LFP power, measured as AUC of PSD) did not significantly differ between infusion groups (all t<0.576) (**Table 1**).

**Table 1:** Baseline values of multi-unit and LFP parameters recorded from the dorsal striatum. *Note*. Baseline values from recordings in the dorsal striatum in the present study are shown alongside values recorded previously from medial prefrontal cortex (Pezze et al., 2014) and ventral hippocampus (McGarrity et al., 2017), using similar recording methods. LFP: local field potential, AUC: Area under the Curve, PSD: power spectral density.

| Parameter | Prospective infusion group | Striatum, present study |  | Medial prefrontal cortex, Pezze et al. (2014) |  | Ventral hippocampus, McGarrity et al. (2017) |  |
| --- | --- | --- | --- | --- | --- | --- | --- |
|  |  | Mean | SEM | Mean | SD | Mean | SEM |
| Mean spike frequency (1/s) | Saline | 10.74 | 4.40 | 17.5 | 5.9 | 42.1 | 11.8 |
|  | Picrotoxin (150 ng) |  |  | 15.6 | 4.3 |  |  |
|  | Picrotoxin (300 ng) | 9.76 | 2.29 | 39.1 | 14.3 | 35.4 | 3.4 |
| Bursts per minute | Saline | 20.24 | 11.78 | 26.48 | 11.54 | 265.5 | 75.5 |
|  | Picrotoxin (150 ng) |  |  | 23.48 | 7.72 |  |  |
|  | Picrotoxin (300 ng) | 16.13 | 4.41 | 47.32 | 13.56 | 251.11 | 27.7 |
| Percentage of spikes fired within bursts | Saline | 38.38 | 7.11 | 69.3 | 4.1 | 28.8 | 3.9 |
|  | Picrotoxin (150 ng) |  |  | 72.3 | 4.6 |  |  |
|  | Picrotoxin (300 ng) | 43.18 | 4.90 | 52.8 | 7.8 | 25.6 | 1.6 |
| Mean burst duration (s) | Saline | 0.30 | 0.09 | 0.23 | 0.03 | 0.0101 | 0.0005 |
|  | Picrotoxin (150 ng) |  |  | 0.23 | 0.03 |  |  |
|  | Picrotoxin (300 ng) | 0.30 | 0.04 | 0.19 | 0.04 | 0.0092 | 0.0002 |
| Mean spikes in burst | Saline | 16.40 | 2.20 |  |  |  |  |
|  | Picrotoxin (150 ng) |  |  |  |  |  |  |
|  | Picrotoxin (300 ng) | 16.85 | 1.82 |  |  |  |  |
| Mean spike frequency in burst (1/s) | Saline | 106.68 | 32.05 | 223.3 | 33.3 | 381.3 | 6.1 |
|  | Picrotoxin (150 ng) |  |  | 218.5 | 22.0 |  |  |
|  | Picrotoxin (300 ng) | 102.12 | 22.39 | 199.9 | 34.6 | 401.8 | 4.9 |
| Mean peak spike frequency in burst (1/s) | Saline | 484.60 | 49.60 |  |  |  |  |
|  | Picrotoxin (150 ng) |  |  |  |  |  |  |
|  | Picrotoxin (300 ng) | 501.59 | 46.02 |  |  |  |  |
| Mean interburst interval (s) | Saline | 12.47 | 4.41 | 6.8 | 2.3 | 6.6 | 1.7 |
|  | Picrotoxin (150 ng) |  |  | 5.8 | 0.97 |  |  |
|  | Picrotoxin (300 ng) | 10.41 | 2.59 | 5.9 | 2.1 | 4.4 | 0.8 |
| Mean burst surprise | Saline | 13.61 | 2.32 |  |  |  |  |
|  | Picrotoxin (150 ng) |  |  |  |  |  |  |
|  | Picrotoxin (300 ng) | 12.27 | 1.46 |  |  |  |  |
| Mean LFP AUC of PSD ($\mu V^2$) | Saline | 0.020 | 0.005 | 0.015 | 0.004 | 0.0411 | 0.0035 |
|  | Picrotoxin (150 ng) |  |  | 0.013 | 0.002 |  |  |
|  | Picrotoxin (300 ng) | 0.020 | 0.004 | 0.024 | 0.006 | 0.0303 | 0.0025 |

We also compared baseline values from our present study in the striatum to baseline values recorded in our previous studies in the mPFC (Pezze et al., 2014) and ventral hippocampus (McGarrity et al., 2017) (**Table 1**). Recordings were made using similar methods, although ventral hippocampal bursts were identified based on the interspike interval (in keeping with other studies on hippocampal bursts), whereas bursts in the present study and in the mPFC were identified using the Poisson method. Mean spike frequency (1/s) in the striatum was about a quarter to half as high as in the mPFC and ventral hippocampus (prospective saline groups, 10.7 vs 17.5 vs 42.1; prospective picrotoxin groups, 9.8 vs 39.1 vs 35.4). Bursts per minute were about 10 to 15 times less in the striatum than in the ventral hippocampus (prospective saline groups, 20.24 vs 265.5; prospective picrotoxin groups, 16.13 vs 251.11), however similar to the mPFC. Mean spike frequency in bursts was about two to four times lower in the striatum compared to the mPFC and hippocampus (prospective saline groups, 106.68 vs 223.3 vs 381.3; prospective picrotoxin groups, 102.12 vs 199.9 vs 401.8). Mean interburst interval was about double in the striatum compared to the mPFC and hippocampus (prospective saline groups, 12.47 vs 6.8 vs 6.6; prospective picrotoxin groups, 10.41 vs 5.9 vs 4.4). Overall, this suggests that baseline burst firing under anesthesia is less intense in the striatum compared to the mPFC and ventral hippocampus.

#### 3.2.3. Striatal picrotoxin infusion enhanced burst firing in the vicinity of the infusion site

Percentage of spikes in bursts (**Figure 3a**), mean spike frequency in bursts (**Figure 3b**) and mean peak spike frequency in bursts (**Figure 3c**) were significantly increased by striatal picrotoxin compared to saline infusion. This was supported by ANOVAs of these parameters showing significant interactions of drug infusion x 5 min block (all F > 2.897, all p < 0.044). In contrast, the mean interburst interval was slightly decreased by striatal picrotoxin compared to saline infusion: more specifically, increase of the interburst interval following saline infusion, probably reflecting a baseline drift of this measure, was not evident following picrotoxin infusion (interaction drug infusion x 5 min block, F_(17,204)_ = 1.840, p = 0.025) (**Figure 3d**). The number of bursts was also increased in the picrotoxin compared to the saline group, which was mainly evident after the picrotoxin infusion, but not during pre-infusion baseline recordings (**Figure 3e**). However, the ANOVA only revealed a main effect of group across pre- and post-infusion 5 min blocks (F_(1,12)_ = 5.193, p = 0.042), but no interaction (F_(17,204)_ = 1.422, p = 0.129) or main effect involving 5 min blocks (F_(17,204)_ = 1.459, p = 0.113). Other multi-unit parameters, including mean burst duration (**Figure 4a**), mean spikes in bursts (**Figure 4b**), mean spike frequency (**Figure 4c**) and mean burst surprise (**Figure 4d**) showed no significant differences between infusion groups (all interactions: F < 1.430 and p > 0.125). For mean spikes in bursts and mean spike frequency, there was an overall effect of 5 min block (both F_(17,204)_ > 1.80, p < 0.046), and there was a trend for such an effect for burst surprise (F_(17,204)_ = 1.567, p < 0.076), reflecting overall changes of these measures across the recording session. In line with the marked LFP spike-wave discharges that were evident in most rats following striatal picrotoxin infusion (see above and **Figure 2b**), overall LFP power, as measured by the AUC of PSD, was numerically increased 2-3 times following striatal picrotoxin compared to saline infusion (**Figure 4e**). However, this difference was not significant (main effect and interaction involving drug infusion, F < 2.7, p > 0.12). This may have partly reflected that the negative spikes of the marked LFP spike-wave discharges were clipped because they were out of our recording range (see **Figure 2b**).

**Figure 3:**
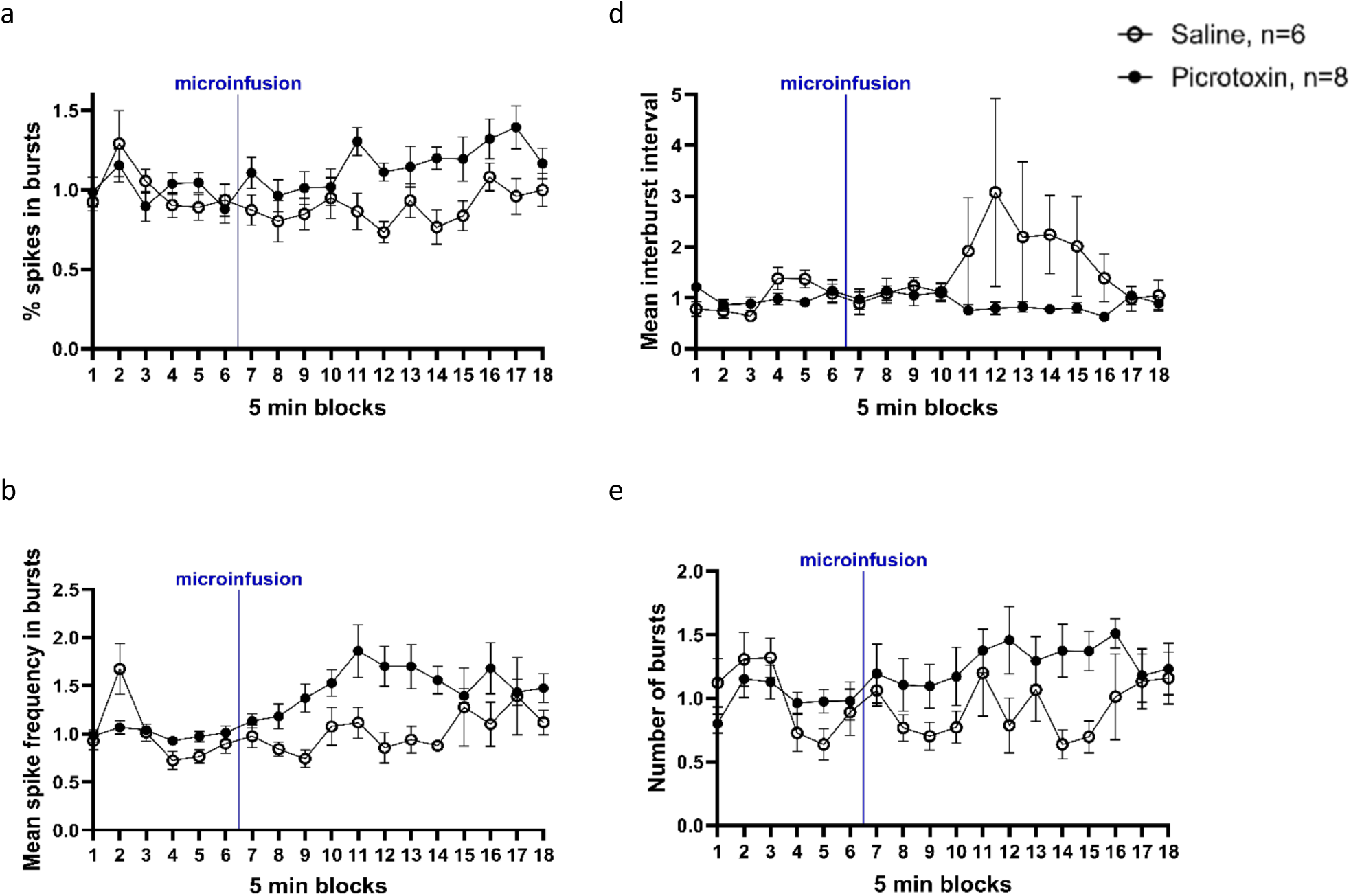
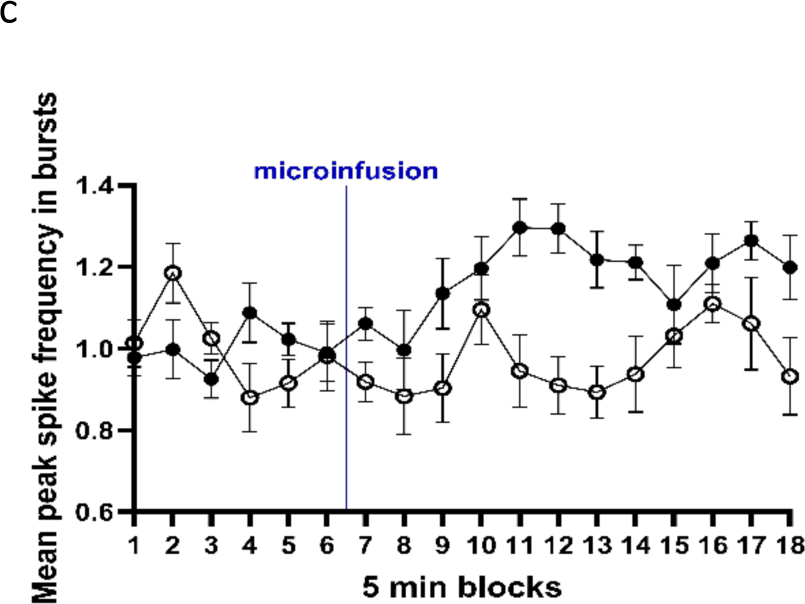
Striatal picrotoxin enhanced multi-unit burst firing parameters in the vicinity of the infusion site. *Note.* Time courses of (a) percentage of spikes in bursts, (b) mean spike frequency in bursts, (c) mean peak spike frequency in bursts, (d) mean interburst interval and (e) number of bursts. Values are normalized to the average of the six baseline 5 min blocks and are presented in mean ± SEM. The vertical blue line after block 6 indicates the time of the striatal saline and picrotoxin infusion.

**Figure 4:**
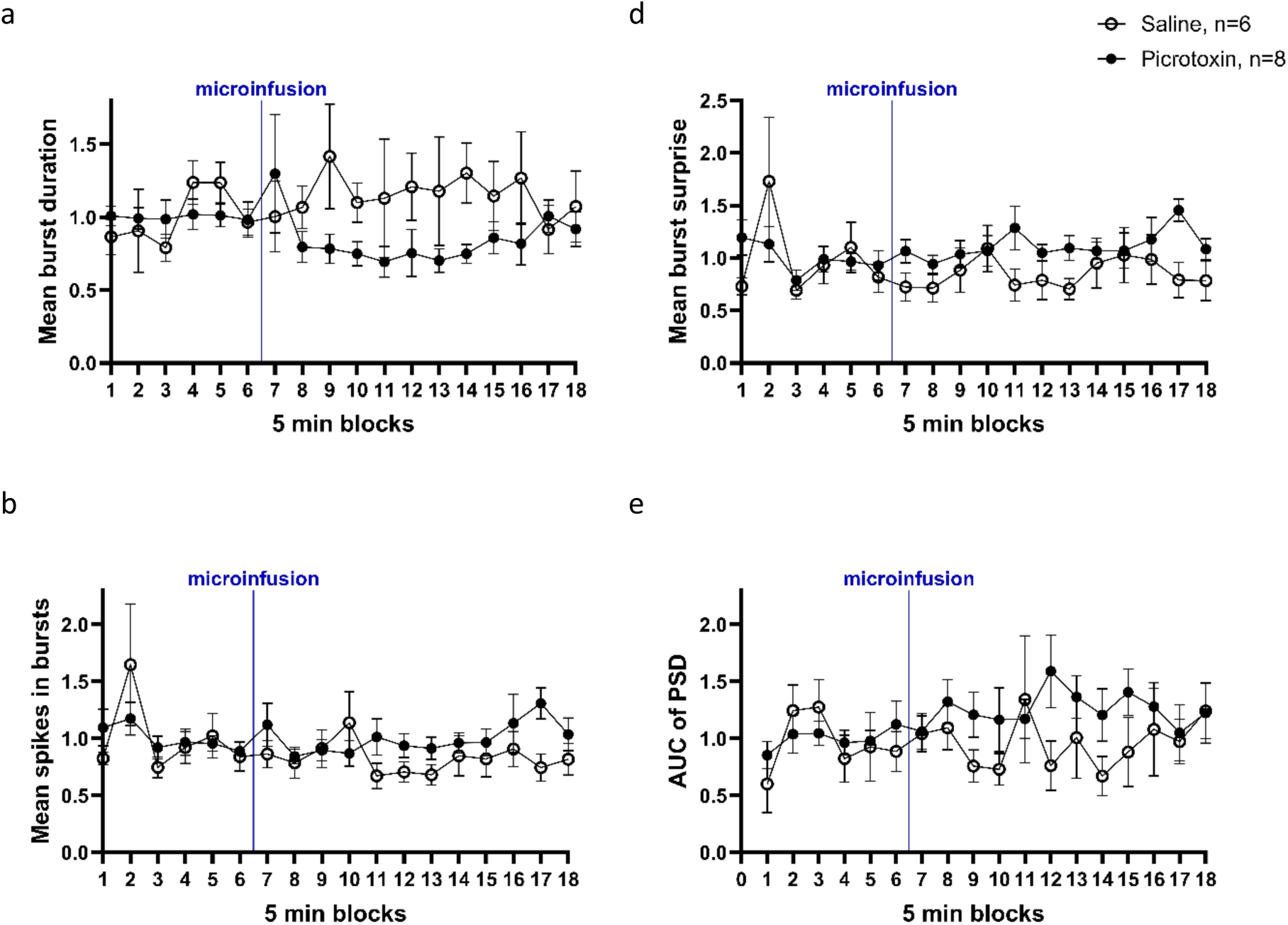
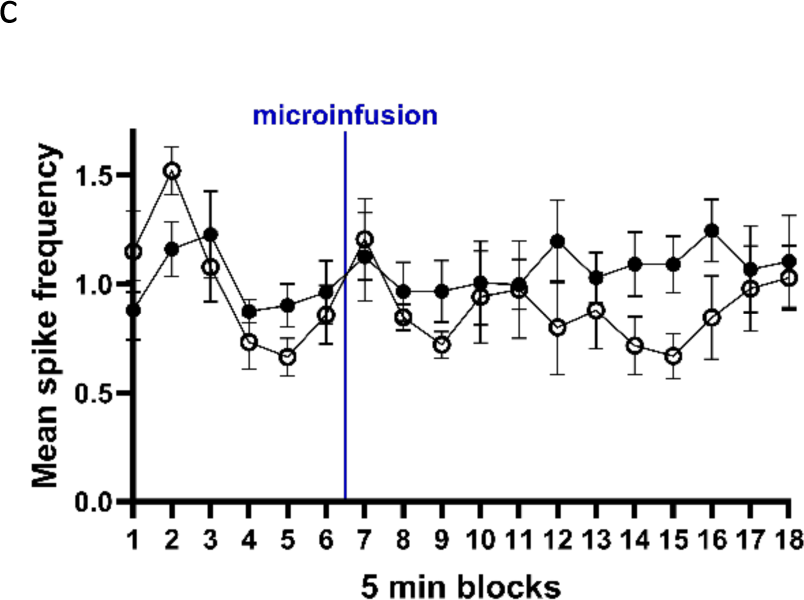
Neural activity parameters in the vicinity of the infusion site that were not affected by striatal picrotoxin. *Note.* Time courses of (a) mean burst duration, (b) mean spikes in bursts, (c) mean spike frequency, (d) mean burst surprise and (e) overall LFP power, measured as Area under the Curve (AUC) of power spectral density (PSD). Values are normalized to the average of the six baseline 5 min blocks and are presented as mean ± SEM. The vertical blue line after block 6 indicates the time of the striatal saline and picrotoxin infusion.

### 3.3. Study 2: Description and time-course of tic-like movements following striatal disinhibition

#### 3.3.1 Description of tic-like movements

Unilateral striatal picrotoxin infusions caused tic-like movements of the contralateral forelimb in all eight rats. Tic-like movements varied in intensity throughout a session, between rats and between infusion days. Some rats started with subtle tic-like movements that became more intense with time. In a typical tic-like movement observed in all rats, the rat lifted its left forelimb, thereby causing its head and torso to rotate to the right around the body’s long axis, before returning the left forelimb, head and torso back into its starting position (**Video 1, Figure 5a**). Across the three picrotoxin infusions (1st and 2nd infusion (day 2 and 6), 300 ng/0.5 µl; 3rd infusion (day 8), 200 ng/0.5 µl), only one out of eight rats showed no tic-like movements on one occasion (following the 2^nd^ infusion, 300 ng), i.e. overall 23 out of a total of 24 picrotoxin infusions evoked tic-like movements.

**Figure 5:**
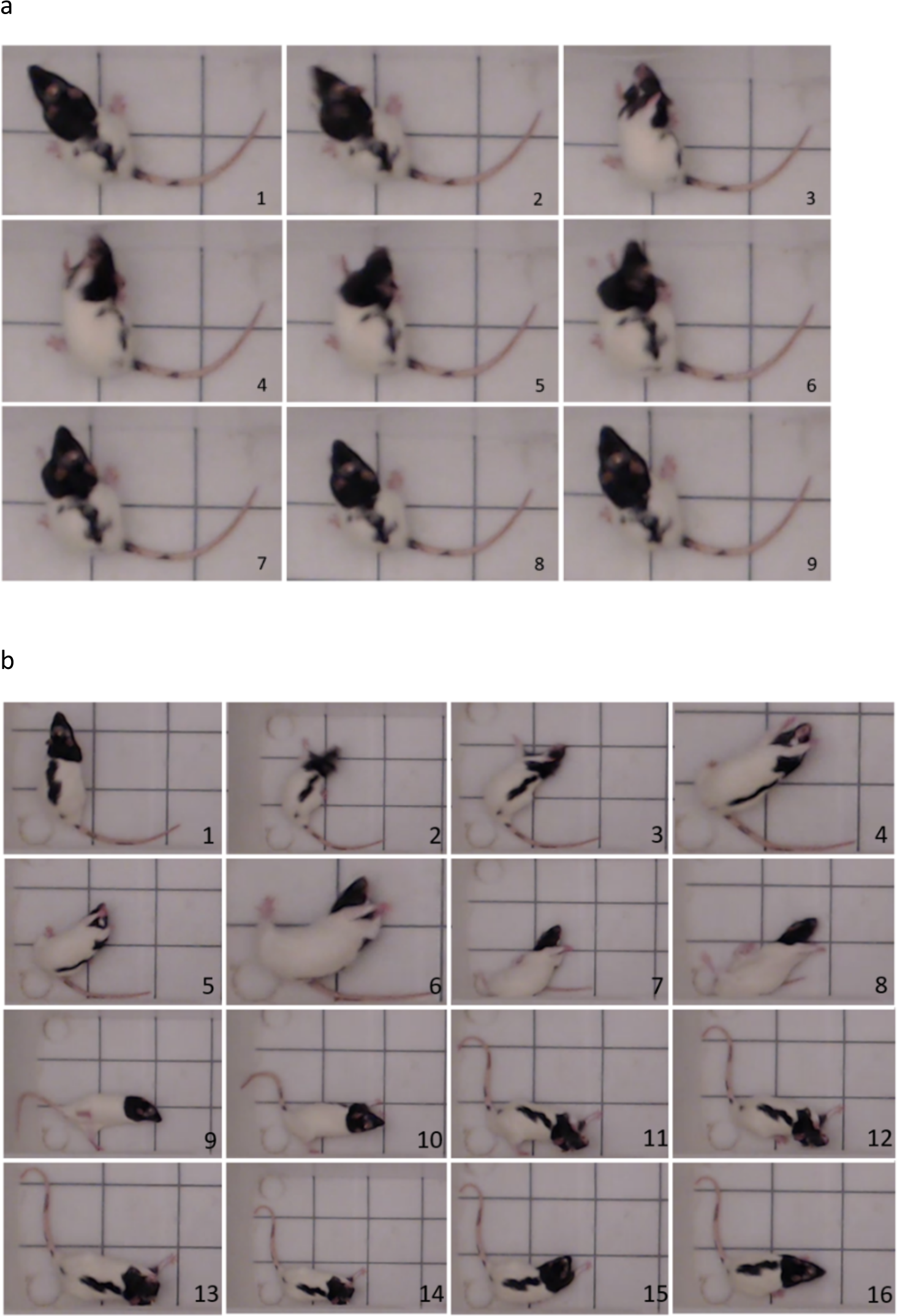
Characteristics of tic-like movements caused by striatal disinhibition. *Note.* Screenshots from video recording of tic-like movements. (a) An example of common tic-like movements that involved lifting the contralateral (left) forelimb, rotation of the head and torso and return to normal body posture. Screenshots were taken from the recording of a rat after picrotoxin infusion (300 ng/0.5 µl) on the second picrotoxin infusion day (day 6). (b) A more pronounced forelimb movement that lasted for several seconds and led to a whole rotation of the whole body around its long axis. Screenshots were taken from the recordings of a rat after picrotoxin infusion (300 ng/0.5 µl), on the first infusion day (day 2).

There were also some more pronounced tic-like movements that lasted for several seconds and led to a whole rotation of the whole body around its long axis (**Video 2, Figure 5b**). This movement started from a normal body posture, with all four limbs placed on the floor (Figure 5b, picture 1), then the rat’s head started rotating around its long axis to the right side and the left forelimb moved forcefully backwards, twisting the rat’s upper body (pictures 2-3). The right forelimb was stretched out, while the left forelimb moved back and forth (pictures 2-8). The rat’s head turned around the long axis, such that the dorsal part of the head came to face the floor and the rat’s mouth came to face upwards (picture 4). With the hind limbs still planted on the ground, the whole body started following the rotational movement of the head (picture 5), such that chest and abdomen came to face upwards, with the head twisted slightly to the right (picture 6). Then, the left hindlimb swung from left to right and the right hindlimb and tail moved along (pictures 6-9), such that the abdomen and mouth came to face toward the floor and the back came to face upward again (picture 10). The rat then stretched the right forelimb out again on the floor and continued to move the left forelimb back and forth with high frequency while the head was twisted to the right (picture 11). The head started moving with the left forelimb, appearing as though the rat was licking its left paw (pictures 13-15). The amplitude of the left forelimb movements reduced, and the head moved to its normal position (picture 15), before the right forelimb moved closer to the body and the rat resumed its normal body position and continued with normal behavior (picture 16).

#### 3.3.2 Time course of tic-like movements

Individual tic-like movements were counted in 5 min blocks. Following striatal disinhibition by picrotoxin infusion, rats started showing tic-like movements within 5 min (**Figure 6**). The number of tic-like movements peaked from about 5 min to 30 min after infusion. All three picrotoxin infusions (300 ng on days 2 and 6, 200 ng on day 8) caused a similar number of tic-like movements across the 12 5 min blocks following infusion. A two-way repeated measures ANOVA revealed no significant difference in the number of tic-like movements (F_(2,14)_ = 1.349, p = 0.291) and no significant interaction between infusion (1^st^, 2^nd^, or 3^rd^) and 5 min block (F_(22,154)_ < 1). There was only a significant main effect of 5 min block (F_(11,77)_ = 10.665, p < 0.001). Following saline infusion (day 4, data not shown), no tic-like movements were observed, nor any other changes in behavior. The mean time to first tic-like movements after striatal picrotoxin infusion and placement of the rat in the box (i.e., the ‘onset’ time) and the duration of tic-like movements did not differ between the three picrotoxin infusions (day 2, 6 and 8) (onset, in s: first infusion, mean = 199; second infusion, mean = 165; third infusion, mean = 157; F_(2,12)_ < 1; duration, in s: first infusion, mean = 1991; second infusion, mean = 3335; third infusion, mean = 2182; F_(2,12)_ = 1.860, p = 0.198; n=7 rats).

**Figure 6:**
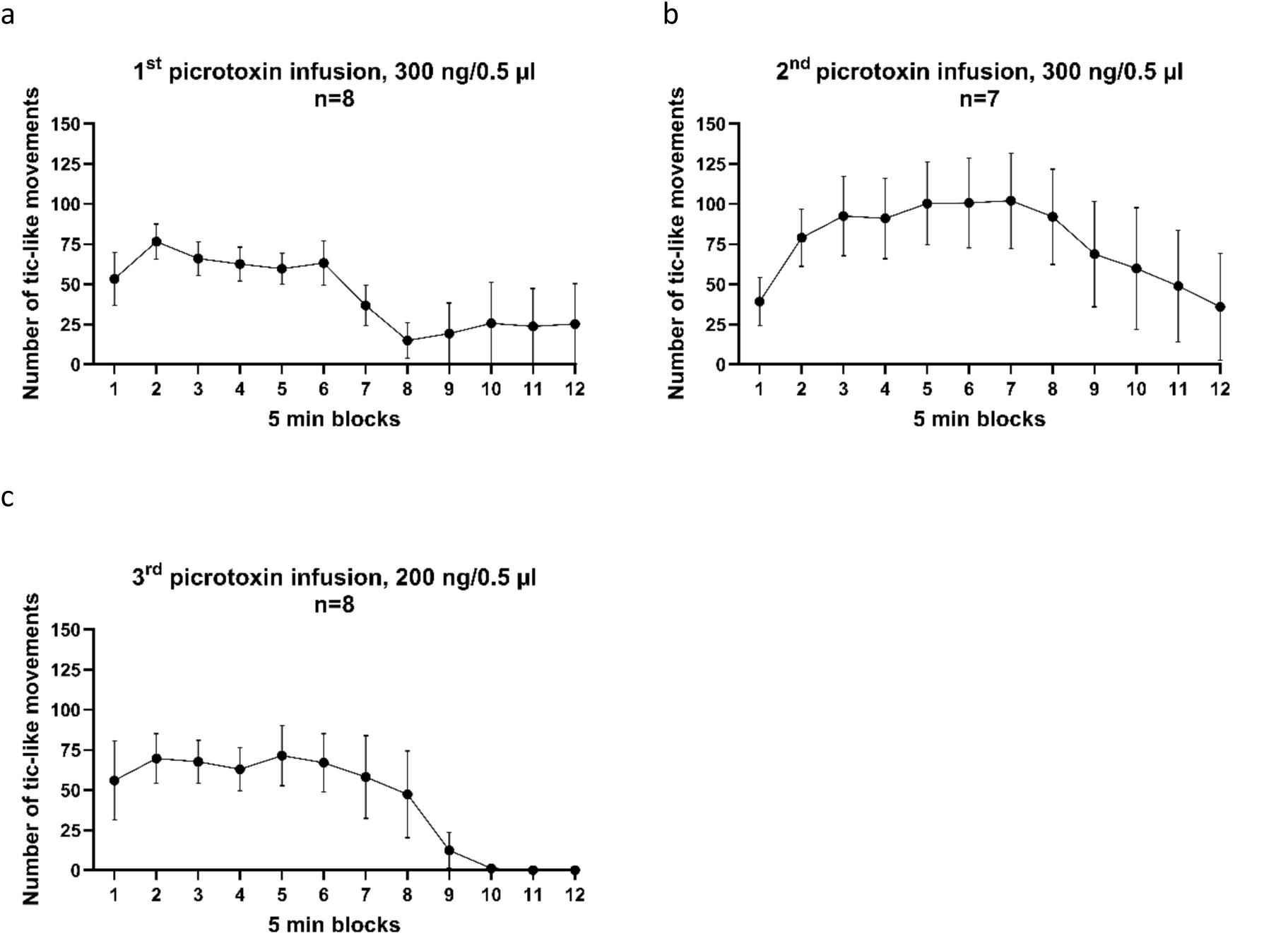
Time course of tic-like movements following striatal disinhibition. *Note.* Tic-like movements (mean ± SEM) are shown in 5 min blocks, following striatal disinhibition by 300 ng (first and second infusion) or 200 ng (third infusion) of picrotoxin in 0.5 µl saline. Regardless of the infusion and picrotoxin dose, tic-like movements peaked from around 5 min (block 1) to about 30 min (block 6). One out of eight rats showed no tic-like movements following the 2nd picrotoxin in 0.5 µl saline infusion and, therefore, the means and SEM were only calculated from n=7 rats.

#### 3.3.3 Separate analysis of tic-like movements involving whole-body rotation

Although most rats showed tic-like movement involving whole-body rotation at some point following striatal disinhibition, these were substantially less frequent than the tic-like movements limited to lifting of the contralateral forelimb and a partial rotation of head and torso. The frequency of tic-like movements involving whole-body rotation varied substantially between individual rats and individual picrotoxin infusions. After the first picrotoxin infusion, 6 out of 8 rats showed some pronounced tic-like movements that involved a whole rotation of the whole body around its long axis, whereas 5 rats showed such movements after the second and third infusions. One rat did not show any tic-like movement involving a whole-body rotation at any point after any of the three picrotoxin infusions. Out of 24 picrotoxin infusions (8 rats, infused with picrotoxin on 3 occasions), rats did not show any tic-like movements involving whole-body rotation in 8 instances. In 9 instances, the movements occurred between 1 and 5 times. In 2 instances, they were observed between 5 and 10 times. In 5 instances, the movements occurred more than 10 times. The highest number of pronounced tic-like movements involving whole-body rotation that was recorded after any infusion was 53 (first picrotoxin infusion, 300 ng/0.5 µl). Although there was a numerical tendency for more tic-like movements involving whole-body rotations after the first (mean = 14.86, SEM = 7.85) and second (mean = 11.43, SEM = 5.37) picrotoxin infusion, when the higher dose (300 ng/0.5 µl) was used, compared to the third picrotoxin infusion (mean = 1.71, SEM = 0.68), when the lower dose (200 ng/ 0.5 µl) was used, a repeated measures ANOVA did not show a significant difference (F_(2,12)_ = 2.118, p = 0.163).

### 3.4. Study 3: Striatal disinhibition tended to reduce startle, without affecting PPI, and increased open-field locomotor activity and non-ambulatory movements

In the startle/PPI experiments, all rats included in the analysis (n=14) showed tic-like movements following striatal picrotoxin. In the open-field experiment, 12 out of 13 rats included in the analysis showed tic-like movements following picrotoxin infusion.

#### 3.4.1 Striatal disinhibition tended to reduce startle, without affecting PPI

Striatal disinhibition tended to reduce startle during the first test block (pulses 1-10), before habituation led to similarly low startle responses in both infusion conditions (**Figure 7a**). There was no significant main effect of drug infusion on startle (F_(1,13)_ < 1), but a strong trend for an interaction of infusion x pulse-alone trials (F_(2,26)_ = 3.183, p = 0.058), alongside a significant effect of test block (F_(2,26)_ = 5.543, p = 0.010), reflecting overall startle habituation.

**Figure 7:**
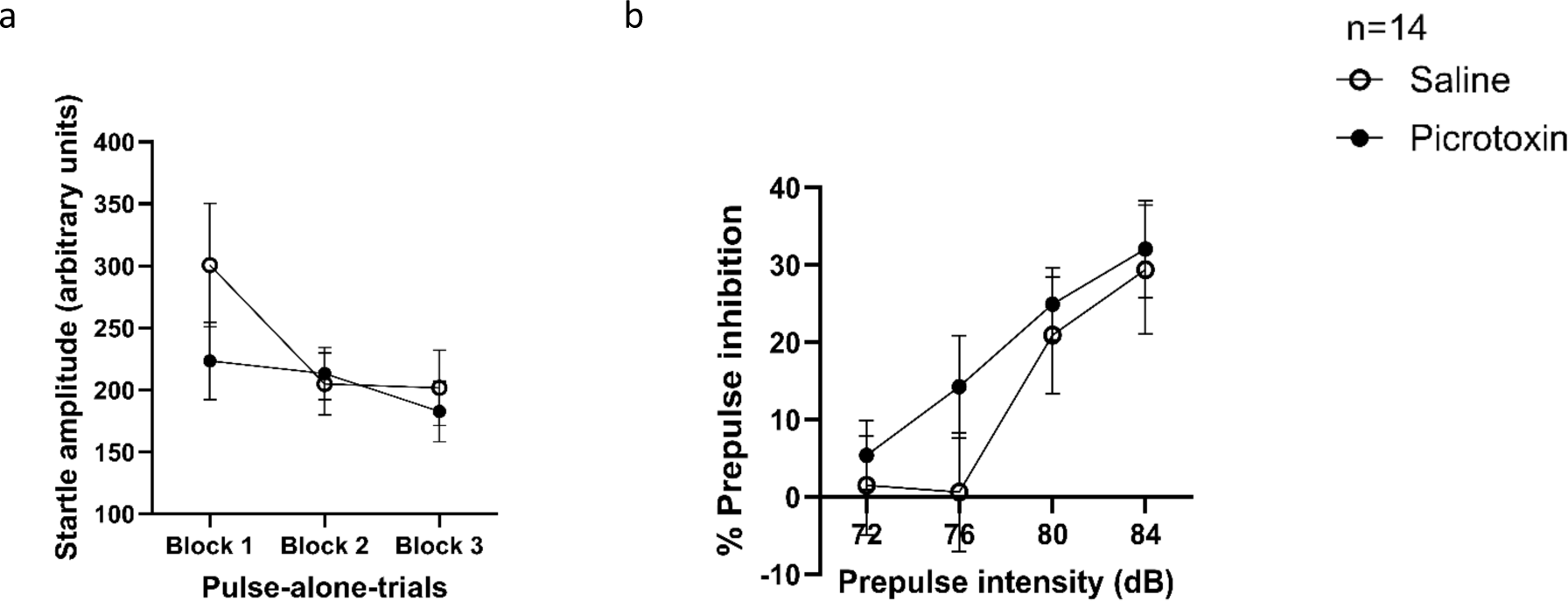
Striatal disinhibition tended to reduce startle, but did not affect PPI. *Note.* (a) Startle response during the three blocks of pulse-alone trials (block 1, trials 1-10; block 2, trials 11-60; block 3, trials 61-65) and (b) percentage of prepulse inhibition (PPI) at the four different prepulse intensities following striatal disinhibition by picrotoxin (300ng/0.5 µl) or striatal saline infusion. Data are shown as mean ± SEM.

Striatal disinhibition did not affect the percentage of PPI (**Figure 7b**). No significant main effect of infusion condition and no significant infusion x prepulse intensity interaction (F < 1, p > 0.44) was found. There was only a significant main effect of prepulse intensity (F_(3,39)_ = 15.452, p < 0.001), with percentage PPI being higher at higher intensities.

#### 3.4.2 Striatal disinhibition increased locomotor activity and non-ambulatory movements

Picrotoxin in the striatum increased locomotor activity (**Figure 8a**). A repeated measures ANOVA revealed a significant effect of drug infusion (F_(1,12)_ = 7.005, p = 0.021), significant effect of time (F_(11,132)_ = 31.498, p < 0.001), reflecting habituation of locomotor behavior, and an interaction (drug infusion x time) that was at trend level (F_(11,132)_ = 1.687, p = 0.083), which reflected that the locomotor hyperactivity caused by striatal disinhibition was most pronounced between 10 and 25 min after start of the test session. Non-ambulatory movement counts were also increased by striatal picrotoxin (**Figure 8b**). This measure increased following striatal picrotoxin, peaking around 20 to 30 min following infusion, before declining to similar levels as in the saline group toward the end of the session. In contrast, following striatal saline, non-ambulatory movements decreased monotonically from the beginning to the end of the session reflecting habituation. These differences between the infusion conditions were supported by a significant interaction of drug infusion x time (F_(11,132)_ = 3.123, p = 0.001), alongside significant effects of drug infusion (F_(1,12)_ = 20.036, p = 0.001) and time (F_(11,132)_ = 8.337, p < 0.001).

**Figure 8:**
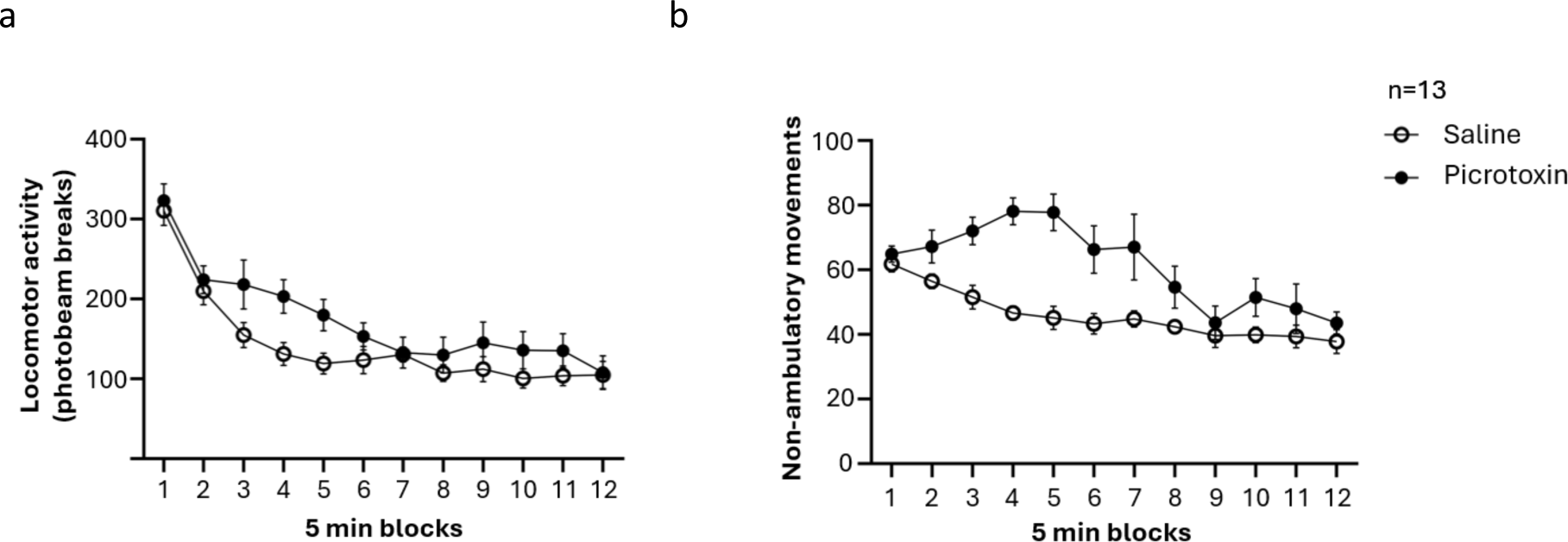
Increased locomotor activity and non-ambulatory movements following striatal disinhibition. *Note.* (a) Locomotor activity and (b) non-ambulatory movements are shown in 5 min blocks, following striatal disinhibition by picrotoxin (300ng/0.5 µl) or striatal saline infusion. Data are shown as mean ± SEM.

During re-baseline testing following infusion days (data not shown), there were no carry-over effects of the picrotoxin infusion, reflected by no significant main effect or interaction involving the infusion condition on the preceding infusion day (all F < 1.309, p > 0.226) for both locomotor activity and non-ambulatory movement count. On re-baseline days, both locomotor and non-ambulatory movements only showed a significant effect of time, reflecting habituation (F > 17.66, p < 0.001).

## 4. Discussion

Right anterior striatal disinhibition by picrotoxin (300 ng/0.5 µl) caused large LFP spike-wave discharges and enhanced multi-unit burst firing in the striatum, without inducing tic-like movements in anaesthetized adult male Lister hooded rats. In freely moving rats, striatal picrotoxin (200 or 300 ng/0.5 µl) reliably caused marked tic-like movements of the left forelimb. Striatal picrotoxin (300 ng/0.5 µl) did not affect PPI, but tended to reduce startle, and increased locomotor activity and non-ambulatory movements in an open-field.

### 4.1. Striatal disinhibition caused large LFP spike-wave discharges and enhanced burst firing: significance for tic-like movements

In anesthetized rats, striatal disinhibition caused large LFP spike-wave discharges, consisting of a single negative spike followed by a positive wave, consistent with previous studies in freely moving and sleeping rats (Israelashvili & Bar-Gad, 2015; Klaus & Plenz, 2016; McCairn & Isoda, 2013; Vinner Harduf et al., 2021). Importantly, under anesthesia, although striatal disinhibition induced marked LFP spike-wave discharges, rats showed no tic-like movements. This shows that striatal LFP spike-wave discharges are not sufficient to cause tic-like movements, supporting a partial dissociation of these phenomena. Such a dissociation has previously been reported in rats and non-human primates with striatal disinhibition (McCairn et al., 2016; Muramatsu et al., 1990; Vinner Harduf et al., 2021) and has been linked to motor cortex inactivation (McCairn et al., 2016; Muramatsu et al., 1990) and lack of entrainment of striatal neuron firing (multi-unit activity) to LFP spikes (Vinner Harduf et al., 2021).

Firing of striatal neurons during the negative LFP spikes was characterized by intense burst firing. Our multi-unit recordings showed that striatal disinhibition specifically enhanced striatal burst firing, reflected by significant increases in mean spike frequency in bursts, mean peak spike frequency in bursts and percentage of spikes fired as part of bursts and by significant decreases of mean interburst interval. This new finding in the striatum is consistent with disinhibition-induced enhancement of burst firing in the mPFC (Pezze et al., 2014) and ventral hippocampus (McGarrity et al., 2017), suggesting that in all these regions GABA-A-receptor mediated inhibition is particularly important to control neural burst firing. Enhanced burst firing in the striatum may contribute to tic-like movements caused by striatal disinhibition in freely moving rats. Previous studies reported enhanced burst firing of striatal projection (medium spiny) neurons in rats with reduced dopaminergic input to the striatum, and such enhanced burst firing has been linked to akinesia (Nisenbaum et al., 1986; Tseng et al., 2001) and catalepsy (Frank & Schmidt, 2003). In rats, where catalepsy was induced by striatal dopamine receptor blockade, striatal burst firing was shown to be strongly associated with tetani of the forelimb muscles (Frank & Schmidt, 2003). Taken together, these findings suggest that enhanced striatal burst firing can contribute to forelimb muscle tonus and, thereby, to both (i) hypokinetic syndromes caused by reduced striatal dopamine transmission, relevant to Parkinson’s disease, and (ii) hyperkinetic syndromes caused by striatal disinhibition, relevant to TS.

### 4.2. Tic-like movements caused by striatal disinhibition

In awake rats, unilateral picrotoxin infusion into the anterior dorsal striatum (300 or 200 ng/0.5 µl) reliably caused tic-like movements of the contralateral forelimb. The time course of tic-like movements quantified in the present study, with a rapid onset following infusion and plateauing of these movements between 5 and 30 min after infusion, is in line with informal observations of the time course of tic-like movements reported following striatal disinhibition (Bronfeld et al., 2013; Patel & Slater, 1987).

Like tics, movements caused by striatal disinhibition look like physiological, natural movements, but occur repetitively and out of context, and are superimposed on ongoing behavior. However, it is impossible to determine if these movements meet other tic criteria, namely if they are preceded by premonitory urges and are actively suppressible (Kurvits et al., 2020), and some authors have referred to similar movements as ‘myoclonus’ (Patel & Slater, 1987; Tarsy et al., 1978) or ‘dyskinesia’ (Crossman, 1987; Muramatsu et al., 1990). For these reasons, we refer to these movements as ‘tic-like’. Typical tic-like movements caused by disinhibition of the right anterior dorsal striatum in the present study consisted of lifting the left forelimb, rotating the head and torso to the right around the body’s long axis, and then returning to the starting position. In TS patients, tics often involve the shoulders, neck and arm (Nilles et al., 2023). In our study, the rats also displayed more intense tic-like movements involving a whole-body rotation around the body’s long axis, alongside back-and-forth movements of the left forelimb. These whole-body rotations, though not naturally observed in freely moving rats, consisted of coordinated movements within the rats’ natural capabilities without inducing harm. Interestingly, tics in individuals with TS can also include body twisting movements and are sometimes referred to as dystonic tics (Baizabal-Carvallo & Jankovic, 2024; Dueck et al., 2009; Jankovic, 1997; Shute et al., 2016). Additionally, alongside tic-like movements, rats exhibited increased locomotor activity, suggesting that the tic-like movements overlay ongoing behavior with minimal interference.

### 4.3. Dorsal striatal disinhibition tended to reduce startle reactivity, but did not affect PPI

Dorsal striatal disinhibition tended to reduce startle reactivity, but did not affect PPI. In rodents with striatal lesions, increased startle reactivity and reduced PPI were observed (Baldan Ramsey et al., 2011; Kodsi & Swerdlow, 1995, 1997). Together, these findings suggest that striatal neural activity negatively modulates startle reactivity. Moreover, although the lesion studies suggest that the striatum is required for PPI, the present study shows that striatal GABAergic inhibition is not, and that PPI is not sensitive to increased striatal activity.

The intact PPI after striatal disinhibition does not support a direct contribution of striatal disinhibition to PPI deficits in TS (Swerdlow, 2013). Therefore, other brain changes in TS may account for the PPI deficits (Koch & Schnitzler, 1997). The reduced startle reactivity observed differs from studies in patients with TS, who showed unchanged (Sachdev et al., 1997; Swerdlow et al., 2001), or heightened startle responses (Gironell et al., 2000; Stell et al., 1995). This difference may stem from compensatory mechanisms in the chronic clinical condition, which could be explored using a chronic striatal disinhibition model.

Swerdlow and colleagues suggested that motor or vocal tics are due to failed automatic gating of sensory information, as reflected by reduced PPI in people with TS (Swerdlow, 2013; Swerdlow & Sutherland, 2005). In line with this suggestion, PPI deficits were reported alongside tic-like movements in transgenic mouse models relevant to TS deficits (Castellan Baldan et al., 2014; Godar et al., 2016; Nasello et al., 2024). However, our findings reveal that acute dorsal striatal disinhibition causes tic-like movements without PPI deficits, indicating that PPI deficits are not necessary for such movements.

### 4.4. Dorsal striatal disinhibition increased locomotor activity and non-ambulatory movement count

Striatal disinhibition-induced locomotor hyperactivity suggests that the dorsal striatum does not only contribute to movements of individual body parts, including tic-like movements, but also to locomotion, consistent with studies linking the striatum to locomotor control (e.g., Jurado-Parras et al., 2020; West et al., 1990; Yoshida et al., 1991). It cannot be excluded that locomotor hyperactivity partly reflects spread of picrotoxin to the ventral striatum. However, the high specificity of tic-like movements and metabolic imaging data (Loayza et al., in preparation) suggest that striatal disinhibition in the present study was mainly limited to the anterior dorsal striatum. Around 60% of patients with TS also have attention deficit hyperactivity disorder (ADHD) (Freeman et al., 2000; Sheppard et al., 1999), which is partly characterized by excess gross motor activity (Sarver et al., 2015). The increased locomotor activity alongside tic-like movements after striatal disinhibition supports that striatal disinhibition may contribute to this common comorbidity of TS.

Striatal disinhibition increased non-ambulatory movements, as measured automatically in the photobeam cages. Interestingly, the time course of this increase (**Figure 8b**) resembled the time course of tic-like movements (**Figure 6**). Both tic-like movements and non-ambulatory movements increased over the first 30 min following infusion before decreasing. The ranges of non-ambulatory movement counts (45 to 80 per 5 min block) and tic-like movements (30 to 75 per 5 min block) were similar. Although these comparisons involve two different rat cohorts, the similar time courses and ranges suggest that non-ambulatory movement counts may partly reflect tic-like movements, and that non-ambulatory movement counts in photo-beam boxes could serve as an automated measure of tic-like movements. However, the high non-ambulatory movement counts in saline-infused rats, which did not exhibit tic-like movements, indicate that these counts also capture non-tic-like movements. Moreover, if every tic-like movement triggered a non-ambulatory count, we would expect higher non-ambulatory counts after picrotoxin infusion than observed in the present study, namely approximately the counts in the saline group plus the number of tic-like movements observed in Study 2.

### 4.5. Conclusions

Our findings suggest that, apart from generating tic-like movements, dorsal striatal disinhibition may contribute to hyperactivity, which is often comorbid with TS. Similar to disinhibition in other brain regions (Bast et al., 2017), striatal disinhibition enhanced striatal burst firing, which may be important for the behavioral effects of striatal disinhibition. Our findings do not support that striatal disinhibition contributes to PPI disruption, which has been associated with TS and suggested to contribute to tic generation.

## Author contributions

JL, SJ and TB designed research.

JL, CT, JR and RGA performed research.

JL analyzed data.

JL and TB wrote the original draft of the paper, with all other authors subsequently contributing to revising the draft; all authors have approved the final content.

## Acknowledgements

We thank the staff of the Biological Service Unit (BSU) for their contributions to the husbandry and welfare of the rats used in the present study. In addition, we thank the School of Life Sciences imaging facility (SLIM) and staff for their contribution to this publication.

